# Structural Basis for Receptor Recognition by Lujo Virus

**DOI:** 10.1101/329482

**Authors:** Hadas Cohen-Dvashi, Itay Kilimnik, Ron Diskin

## Abstract

Lujo virus (LUJV) has emerged as a novel and highly fatal human pathogen. Despite its membership among the *Arenaviridae*, LUJV does not classify with the known Old and New World groups of that viral family. Likewise, LUJV was recently found to use neuropilin-2 (NRP2) as a cellular receptor instead of the canonical α-dystroglycan (α-DG) or transferrin receptor 1 (TfR1) utilized by Old World (OW) and New World (NW) arenaviruses, respectively. The emergence of a deadly new pathogen into human populations using an unprecedented entry route raises many questions regarding the mechanism of cell recognition and the risk that Arenaviruses are further diversifying their infection strategies. To provide the basis for combating LUJV in particular, and to increase our general understanding of the molecular changes that accompany an evolutionary switch to a new receptor for Arenaviruses, we used X-ray crystallography to reveal how the GP1 receptor-binding domain of LUJV (LUJV_GP1_) recognizes NRP2. Our structural data imply that LUJV is evolutionary closer to OW than to NW arenaviruses. Structural analysis supported by experimental validation further suggests that NRP2 recognition is metal ion dependent and that the complete NRP2 binding is formed in the context of the trimeric spike. Taken together, our data provide the mechanism for the cell attachment step of LUJV, the evolutionary relationship between the GP1 domain of this novel pathogen and other arenaviruses, and indispensable information for combating LUJV.

## Introduction

The *Arenaviridae* family of viruses includes viruses that reside in rodent hosts (mammarenaviruses), producing chronic, mostly asymptomatic infections^1^. Through the consumption of contaminated foods or by inhaling aerosolized feces, several mammarenaviruses can infect humans and cause acute diseases, including hemorrhagic fevers with high morbidity and fatality rates^1,2^. These enveloped, bi-segmented RNA viruses are genetically and geographically divided to the OW mammarenaviruses, endemic to West Africa, and the NW mammarenaviruses, endemic to South and North America^3^. OW mammarenaviruses, including the notorious Lassa virus (LASV), utilize α-DG as a cell entry receptor^4,5^. Most of the NW mammarenaviruses, including the pathogenic Junín, Machupo, Guanarito, and Sabiá viruses, utilize TfR1 as an entry receptor^6,7^. Receptor recognition is achieved by the GP1 receptor-binding domains, which are part of the class-I trimeric spike complexes on the viral surfaces^8^. Following viral internalization into an endocytic compartment of the cell, the spike complexes mediate pH-dependent fusion of the viral and intracellular membranes to deliver the viral genomes into the cell cytoplasm^9^. The GP1 domains of both OW and NW mammarenaviruses adopt the same fold^10-14^, but isolated GP1 domains from OW mammarenaviruses show substantial conformational differences compared with their trimer-associated counterparts^11,12,15^. During cell entry, LASV dissociates from its α-DG receptor and binds lysosomal-associated membrane protein 1 (LAMP1)^16^. We have found that the binding to LAMP1 triggers membrane fusion by the spike complex of LASV^17^, and that utilizing LAMP1 is a unique mechanism for LASV^12^. In the case of LASV, the altered conformation that GP1 adopts is needed for LAMP1 binding^11^ and may have other uncharacterized functions.

Recently, LUJV was identified as a novel pathogenic mammarenavirus during a limited but highly fatal outbreak in Southern Africa^18^. Although LUJV has emerged in Africa, which serves as the ecological niche for most of the OW mammarenaviruses, phylogenetic analysis indicated that LUJV is distinct from the OW and NW mammarenaviruses^18^. A genetic screen revealed that LUJV utilizes NRP2 as a cell entry receptor and that the tetraspanin CD63 must also be present for productive cell entry^19^. This study also showed that the binding of LUJV to NRP2 is reduced in acidic conditions^19^, which is reminiscent of the LAMP1 switching mechanism in LASV. To uncover the structure of the receptor-binding module of LUJV, to reveal how LUJV recognizes NRP2, and to elucidate the evolutionary relationship between LUJV and the OW and NW mammarenaviruses, we studied the structure of LUJV_GP1_ in complex with NRP2 using X-ray crystallography. Analysis of the structure provided the molecular mechanism that allows LUJV_GP1_ to bind NRP2 and uncovered the involvement of a metal ion in mediating this recognition. Structural analysis further suggests the formation of a quaternary binding site for NRP2 in the context of the trimeric spike. The structure of LUJV_GP1_ implies that LUJV is evolutionarily closer to OW than to NW mammarenaviruses, but unlike the OW viruses LUJV_GP1_ seems to maintain its trimer-associated conformation as an isolated domain, an insight that will be important for devising strategies to combat LUJV.

## Results and discussion

### Determining the structure of LUJVGP1 in complex with NRP2

To reveal how LUJV recognizes NRP2, we co-expressed 6xHis-tagged versions of LUJV_GP1_ (residues 74-199) and the first CUB domain of NRP2 (residues 27-144) as secreted proteins using insect cells. The LUJV_GP1_/NRP2 complex was purified by immobilized metal affinity chromatography followed by size exclusion chromatography. The LUJV_GP1_/NRP2 complex formed thin needle-like crystals from which we were able to collect X-ray diffraction data to 2.44 Å resolution (Supplementary Table 1). Using the available structure of NRP2^20^ (PDB: 2QQK) as a search model for molecular replacement (MR) in Phaser^21^, we were able to find 2 copies of the first CUB domain in the asymmetric unit. In contrast, none of the available GP1 structures from mammarenaviruses or homology models based on their structures were able to provide a MR solution for LUJV_GP1_. We hence used phases from the partial solution of NRP2 to trace LUJVGP1 into electron density maps. Initial tracing was done with Phenix.AutoBuild^22^, followed by manual building of the model into density-modified maps using Coot^23^. The final model consists of residues 87-196 for LUJV_GP1_ and residues 24-143 for NRP2, with two copies of the complete complex that relate to each other with translational non-crystallographic symmetry (Fig. 1a). Electron density for residues 74-86 of LUJV_GP1_ and residues 144-145 of NRP2 was missing and hence these residues were not modeled.

**Figure 1:**
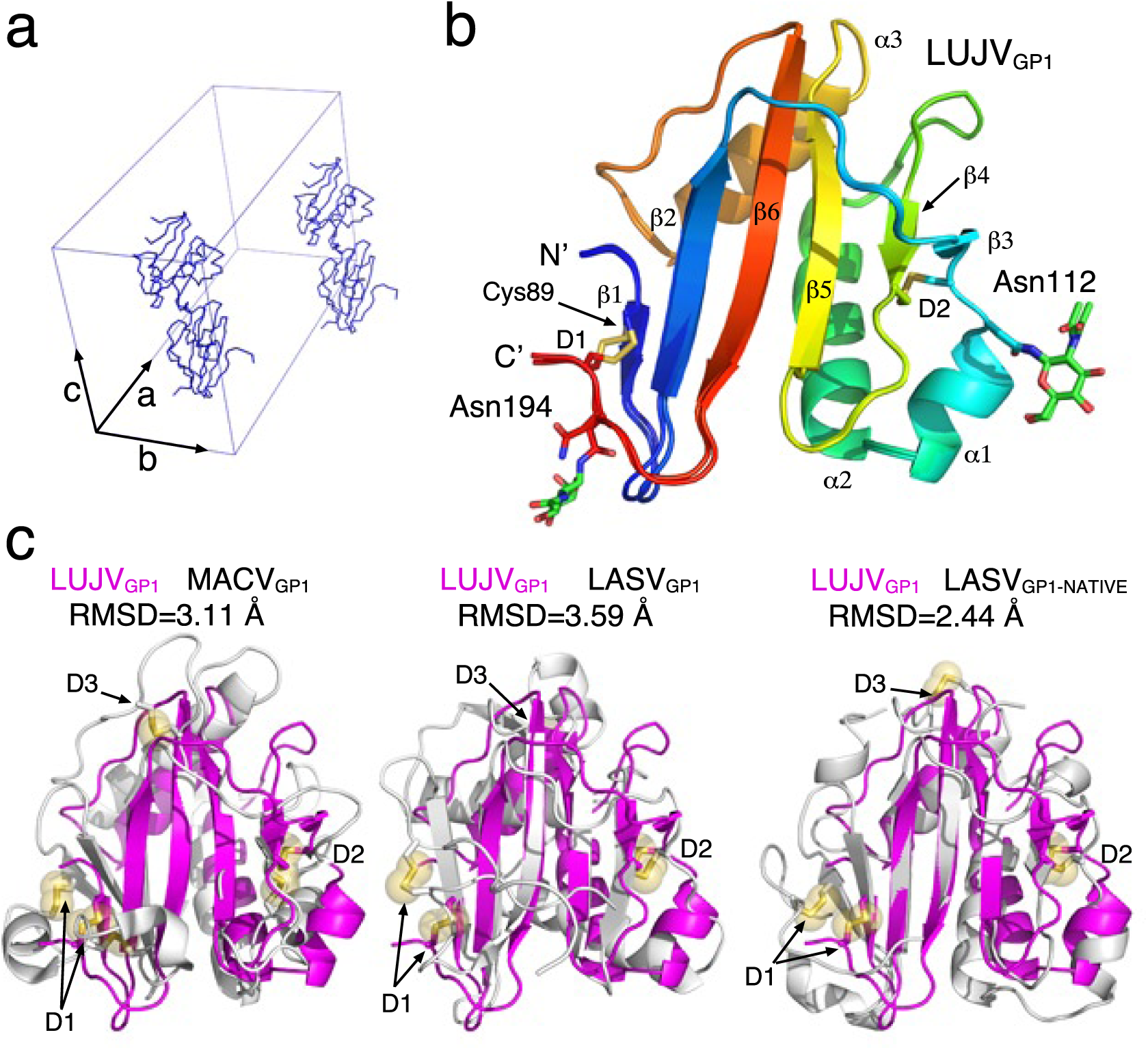
Crystal structure of the GP1 receptor binding domain from LUJV. **a**, The asymmetric unit. The orthorhombic unit cell is shown with blue edges. The two copies of the LUJV_GP1_/NRP2 complex that make the asymmetric unit are shown using a Cα trace. The two copies relate to each other by fractional translations of 0.5, 0.447, and 0.149 along the ‘a’, ‘b’, and ‘c’ axes, respectively. **b**, Overall structure of LUJV_GP1_. Ribbon diagram showing the two copies of LUJV_GP1_ superimposed. The color of the ribbons changes along the primary structure of the protein (N’ terminus, blue; C’ terminus, red). Secondary structure elements, disulfide bonds (D1 & D2), chain termini, and the glycosylation sites that were observed and modeled are annotated. **c**, Superimpositions of LUJV_GP1_ with representative GP1 structures. GP1 domains of Machupo virus (MACV)^10^ (PDB: 2WFO), Lassa virus (LASV)^11^ (PDB: 4ZJF), and LASV in the trimer-associated native conformation^15^ (PDB: 5VK2) are shown in gray ribbons in the left, middle, and right panels, respectively. LUJV_GP1_ is shown in magenta ribbon. These alignments was generated based on the Cα atoms of the central β-strands. RMSD was calculated for each pair of structures using TM-align^15^. RMSD calculations were based on 107, 109 and 74 Cα atoms for MACV, LASV, and LASVNATIVE, respectively.

### LUJV_GP1_ adopts a structure that resembles the trimer-associated native conformation of ‘Old World’ mammarenaviruses

Based on its glycoprotein spike complex (GPC) sequence, LUJV does not cluster with OW or NW mammarenaviruses^19^. LUJV further uses a distinct receptor compared to other viruses in this family^19^. It is thus unclear how different is the GP1 receptor-binding domain of LUJV compared to the other viruses from this family. The LUJV_GP1_/NRP2 crystal structure reveals that LUJV_GP1_ has a central β-sheet flanked by loops on one side and helices and loops on the other (Fig. 1b), in a configuration that resembles the structures of GP1s from OW^11,12,15,24^ and NW^10,13,14^ mammarenaviruses. Comparing LUJV_GP1_ to GP1 domains from LASV and Machupo virus (MACV) as representatives of OW and NW viruses reveals interesting structural differences (Fig. 1c). Whereas the central β-sheet is somewhat conserved, with the exceptions of a longer β-strand 6 in LUJV_GP1_ and the absence of an N’ terminal β-strand preceding β1 of LUJV_GP1_ (Fig. 1b and 1c), the relative orientations of the helices with respect to the central β-sheet are not maintained (Supplementary Fig. 2). In addition, the conformations of most of the loops that connect the secondary structure elements greatly differ. A prominent β-hairpin^11^, which contributes to the LAMP1 binding site on LASV_GP1_^12^ but is shared by other OW mammarenaviruses that do not utilize LAMP1 during cell entry^12^, is missing in LUJV_GP1_. Instead, this hairpin is reduced in LUJV_GP1_ to a short loop that connects α-helix 3 with β-strand 6 similarly to GP1 domains from NW mammarenaviruses (Fig. 1b and Supplementary Fig. 1). However, LUJV_GP1_ is missing a disulfide bond (D3) in this region that is found in both OW and NW mammarenaviruses (Fig. 1b and 1c). Despite these differences and the low overall sequence similarity, LUJV_GP1_ clearly belongs to the same fold family as GP1 domains from other arenaviruses.

Interestingly, GP1 domains from OW mammarenaviruses like LASV and Morogoro virus adopt a characteristic altered conformation^11,12^ compared to the trimer-associated native conformation^15^. In the case of LASV, this altered conformation makes it compatible for binding LAMP1^11,17^, and it may further serve for immunological evasion. These conformational changes of LASV_GP1_ involve the rearrangement of termini that disrupt the first β-strand as well as significant rearrangements of the helices (Supplementary Fig. 3). The termini of LASV as well as of all the GP1 structures from other mammarenaviruses solved to date are linked by a characteristic disulfide bond (D1) (Fig. 1b, 1c and Supplementary Fig. 1). In LUJV_GP1_ however, the termini are linked differently by a disulfide between Cys195 at the C’ terminus and Cys89 on β1. Cys89 of LUJV is not located on equivalent region as in GP1 from MACV or LASV (Fig. 1c and Supplementary Fig. 1). In the structure of LASV_GP1_ that was determined in the context of the native trimeric spike complex^15^, the cysteine that corresponds to Cys89 of LUJV is located on the extra β-strand that precedes β-strand 1 of LUJV_GP1_ (Fig. 1c). Likewise, the equivalent disulfide bond in MACV also links the C’ terminus of the protein to an extra β-strand at the N’ terminus. In the current crystal structure, the first 13 residues of LUJV_GP1_, which could potentially have formed such extra β-strand, are disordered. Although the absence of a preceding β-strand may resemble one attribute of the altered LASV_GP1_ conformation when isolated, the overall structure of LUJV_GP1_ is more similar to the trimer-associated conformation of LASV_GP1_ than to isolated LASV_GP1_ or to MACV_GP1_. This similarity is visually evident and further reflected by the lower RMSD (Fig. 1c and Supplementary Fig. 2). This observation implies that LUJV_GP1_ as an isolated domain maintains its trimer-associated native structure.

### LUJV_GP1_ recognizes NRP2 using an intricate network of polar interactions

Two types of interfaces between LUJV_GP1_ and NRP2 are found in the crystal structure (Supplementary Fig. 4). One involves a flat surface on NRP2 and has a combined buried surface area (BSA) of 624 Å^2^(309 and 315 Å^2^on LUJV_GP1_ and NRP2, respectively). The second interface has a combined BSA of 1158 Å^2^ (613 and 545 Å^2^on LUJV_GP1_ and NRP2, respectively) and involves loops from both proteins that together with their side-chains form geometrically compatible interaction surfaces (Fig. 2a). We thus consider the first interface as a crystal contact and the second as the biologically relevant binding interface. On LUJV_GP1_, the short β-strand 3 and its flanking residues, as well as the loop that connects α-helix 2 with β-strand 4 (α2β4 loop), form most of the binding site (Fig. 2a). A major NRP2 determinant recognized by LUJV_GP1_ is a calcium-binding site formed by loops on the receptor surface (Fig. 2b). Though calcium binding sites were already observed in other domains of NRP2^20^, this site on the first CUB domain was not occupied in the previously determined structure (supplementary Fig. 5). The binding of a calcium ion stabilizes the conformations of Asp127 and Glu79 from NRP2 (supplementary Fig. 5) pre-organizing them for interaction with Lys110 of LUJV_GP1_ (Fig. 2b). By coordinating a main-chain carbonyl of Arg130 (Fig. 2b), the calcium ion stabilizes the loop between residues 128-130 of NRP2, which is otherwise mobile (supplementary Fig. 5). The conformation of this loop is important for LUJV_GP1_ binding, as it positions Arg130 of NRP2 to form hydrogen bonds with two main-chain carbonyls from the LUJV_GP1_ α2β4 loop (Fig. 2b). This particular rotamer of Arg130 that LUJVGP1 recognizes is stabilized by a hydrogen bond between the Arg130 guanidino group and the calcium-positioned carboxylic acid of Glu79 (Fig. 2b). In the absence of a calcium ion, the local architecture of NRP2 would be disrupted (supplementary Fig. 5), which will likely prevent binding of LUJV_GP1_. Indeed, the first NRP2-CUB domain fused to an Fc portion of an antibody can stain HEK293 cells that express the LUJV spike complex as long as EGTA is not added to the staining solution (Fig. 2c).

**Figure 2:**
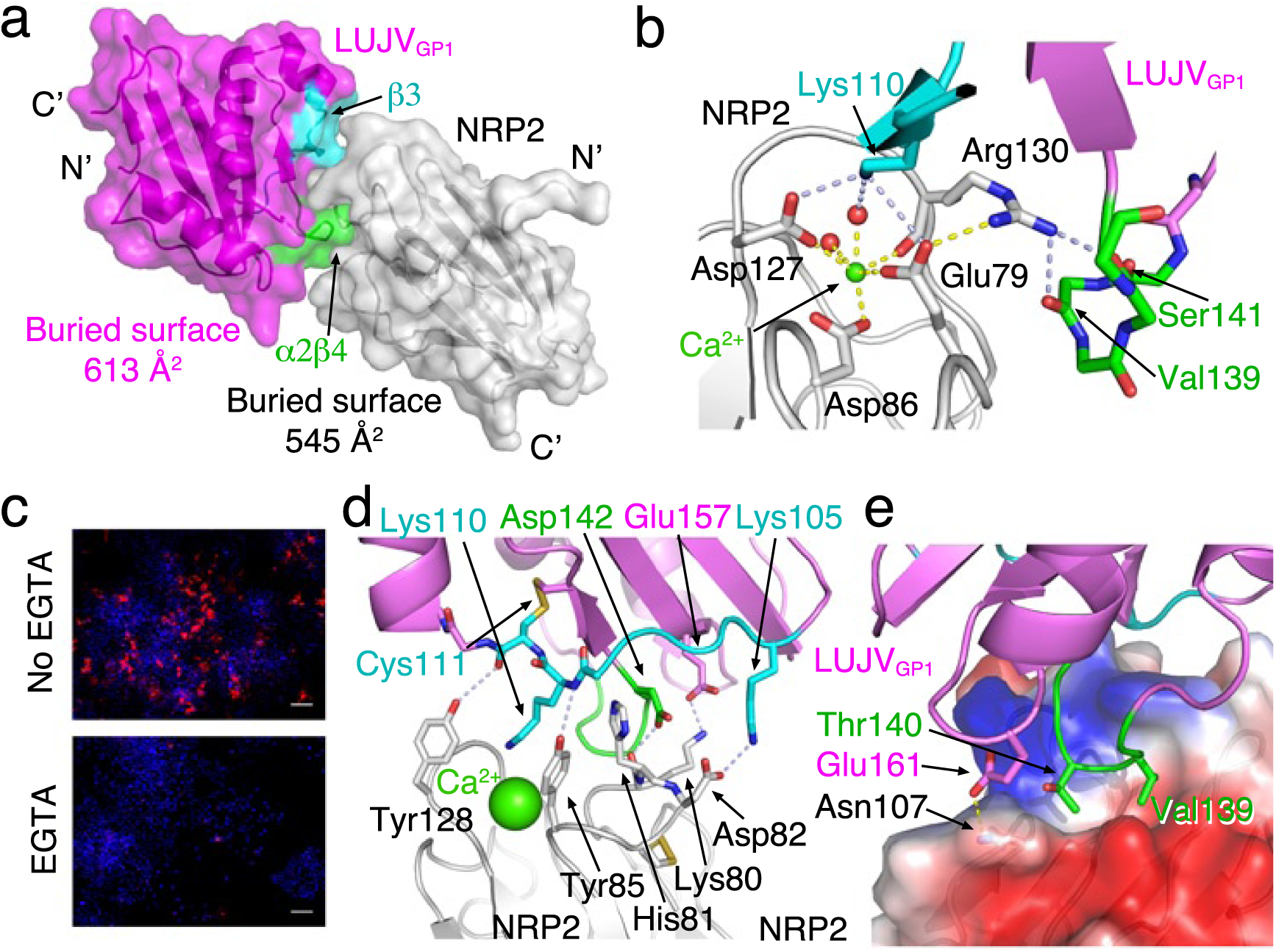
NRP2 recognition by LUJVGP1. **a**, Combined ribbon and surface representations of the LUJV_GP1_/NRP2 complex in magenta and gray for LUJV_GP1_ and NRP2, respectively. The buried surface area on each protein is indicated as well as the locations of the termini. β-strand 3 and the α2β4 loop of LUJV_GP1_ are highlighted in cyan and green. **b**, A calcium coordination site on NRP2 as a central recognition element of LUJV_GP1_. Side-chains and important main-chain elements of this interaction motif are shown as sticks. Polar interactions are marked using either yellow dashed lines for intramolecular interactions or light-blue dashed lines for interactions between LUJV_GP1_ and NRP2. A calcium ion is indicated as a green sphere, and its coordination by three carboxyl groups, one main-chain carbonyl and two water molecules (red spheres) is illustrated. The color scheme of LUJV_GP1_ is the same as in ‘a’. **c**, Calcium is required for LUJV_GP1_/NRP2 interaction. Fluorescence microscopy images of HEK293 cells that were transiently transfected with a plasmid encoding the spike complex of LUJV. The cells were stained with a chimeric protein made of the first CUB domain of NRP2 fused to Fc portion of an IgG1. Bound chimera is visualized using a fluorescently labeled anti-human antibody (red). Nuclei are stained with DAPI (blue). The addition of 5 mM EGTA abolished staining (lower image) as compared to the control (upper image). Size bars indicate 50 μm. **d**, Additional polar interactions between LUJV_GP1_ and NRP2. This panel uses the same color and representation scheme as in ‘b’. The calcium ion is shown using a green space-filling sphere for orientation. **e**, Apolar interaction motif with NRP2. The coloring scheme for LUJV_GP1_ is the same as is ‘a’. NRP2 is shown using a surface representation colored by the local contact electrostatic potential. Blue represents positive potential, red represents negative potential and white represents apolar surfaces.

The loop that stretches from Glu79 to Asp86 of NRP2 bears several charged residues that are also utilized by LUJV_GP1_ for binding. Both Lys80 and Asp82 of NRP2 form salt-bridges with the counter-charged Glu157 and Lys105 of LUJV_GP1_, respectively (Fig. 2d). The main-chain carbonyl group of Lys80 from NRP2 forms a polar interaction (likely a hydrogen bond) with Asp142 of LUJV_GP1_. In addition, Tyr85 and Tyr128 of NRP2 form hydrogen bonds with the main-chain amide group of Lys110 and the main-chain carbonyl group of Cys111 from LUJV_GP1_, respectively. Beside these polar interactions, Val139 and Thr140 from the α2β4 loop of LUJV_GP1_ participate in Van der Waals interactions inside a hydrophobic pocket that is formed by the aliphatic portions of some NRP2 residues like Glu79 and Glu77 (Fig. 2e). Additional hydrophobic interactions are formed by His81 of NRP2 with non-polar atoms on LUJV_GP1_ (Fig. 2d). An adjacent Asn107 on the surface of NRP2 makes yet another hydrogen bond with Glu161 from LUJV_GP1_. Altogether, the interface between LUJV_GP1_ and NRP2 is mostly polar.

### The NRP2 binding site on LUJV_GP1_ spans a region that mediates α-dystroglycan recognition by OW mammarenaviruses

Previous studies provided structural and biochemical data for the recognition of α-DG and TfR1 receptors by OW and NW mammarenaviruses. Structural studies by Abraham *et al.*^25^ illustrated that the TfR1 binding site on GP1 of NW mammarenaviruses is formed on the face of the central β-sheet together with the flanking loops (Fig. 3a). By superimposing LUJV_GP1_ with MACV_GP1_ crystallized in complex with TfR1, it is evident that the NRP2 recognition site comprises a different surface of GP1 (Fig. 3a). In contrast, the NRP2-binding surface of LUJV_GP1_ appears to be close to residues that contribute to α-DG binding in OW mammarenaviruses (Fig. 3b). Previous studies with the α-DG-tropic OW lymphocytic choriomeningitis virus (LCMV) pointed to a few GP1 residues important for binding to its α-DG receptor^26,27^. These residues were later mapped on the structure of the trimeric spike complex of LASV and found to be at its apex near the trimer interface^15^. Interestingly, Tyr150, Asn148, and Ile254 of LASV that contribute for α-DG binding appear at the NRP2 binding site when LASV_GP1_ is superimposed on LUJV_GP1_ (Fig. 3b). Although LUJV and LASV do not share any similar sequences or local loop conformations in this particular region, LUJV has evolved to utilize the same overall region of GP1 for NRP2 recognition as OW mammarenaviruses use for binding α-DG. This observation implies that LUJV is evolutionary closer to the OW than the NW mammarenaviruses, a notion that is further supported by the higher structural similarity of LUJV_GP1_ to LASV_GP1-NATIVE_ compared with MACV_GP1_ (Fig. 1c). In the absence of structural information for α-DG recognition by OW mammarenaviruses, the LUJV_GP1_/NRP2 structure may further hint about additional GP1 regions that have a potential to contribute for binding to α-DG.

**Figure 3:**
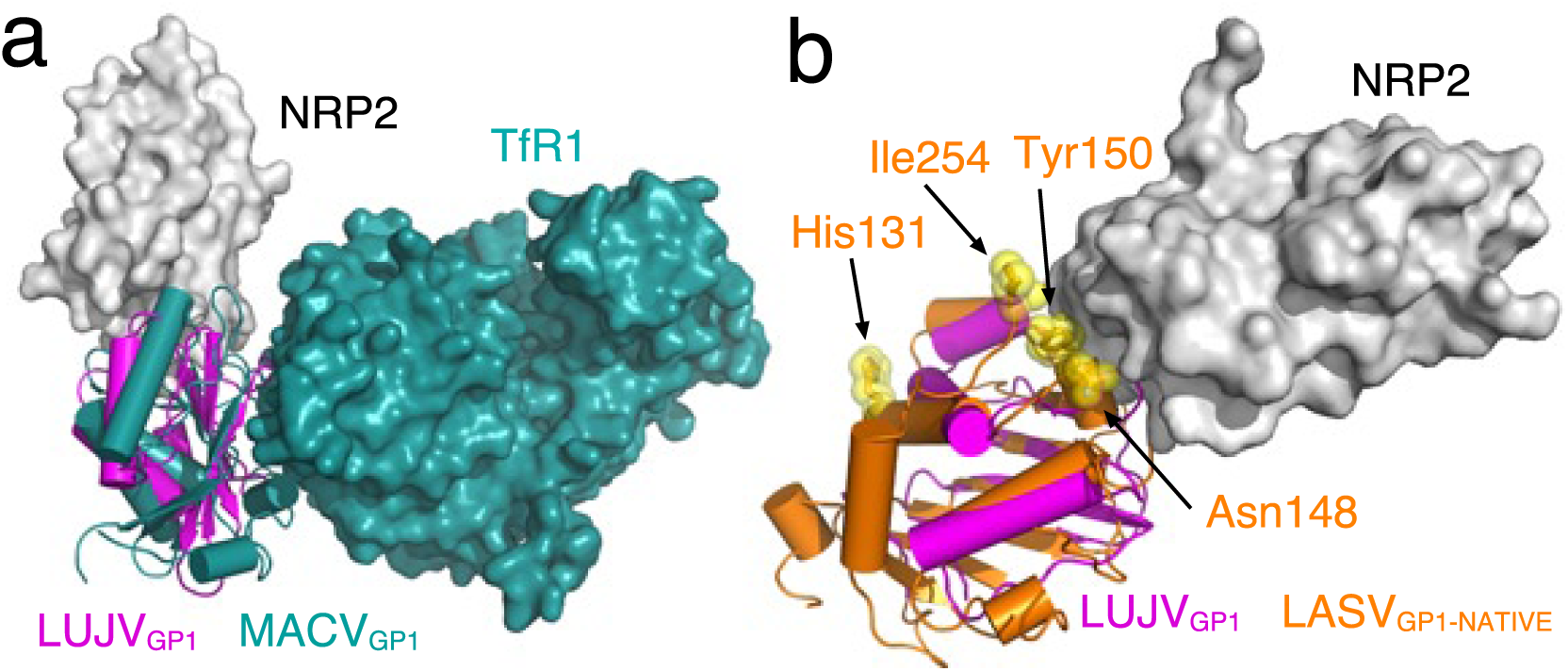
The NRP2 binding site on LUJV_GP1_ is located closer to the α-DG binding site than to the TfR1 binding site. **a**, Superimposition of LUJV_GP1_/NRP2 on the MACVGP1/TfR1 complex^25^ (PDB: 3KAS). The GP1 domains are shown using ribbon representation in magenta and bottle green color for LUJV_GP1_ and MACV_GP1_, respectively. TfR1 and NRP2 are shown using surface representations in bottle green and gray, respectively. There is no overlap between the NRP2 and TfR1 binding sites. **b**, The LUJV_GP1_/NRP2 complex is superimposed on LASV_GP1_ in the trimer-associated conformation (orange). Tyr150, Asn148, and Ile254 of LASV, which participate in α-DG recognition, are highlighted. The highlighted His131 of LASV is also contributing for α-DG recognition by joining with Tyr150, Asn148, and Ile254 of a neighboring LASV_GP1_ in the context of the trimer spike complex.

### NRP2 recognition in the context of the trimeric spike hints for a combined quaternary binding site

Since LUJV_GP1_ superimposes reasonably well with LASV_GP1_ in its trimeric, associated, native conformation (Fig. 1c), we utilized the trimeric structure of the LASV spike complex that was determined by Hastie *et al.*^15^, to gain insights about how NRP2 might be recognized in the context of the trimeric spike. By superimposing the LUJV_GP1_/NRP2 complex on the three copies of LASV_GP1-NATIVE_, we found that NRP2 will bind the apex of the spike complex (Fig. 4a).

**Figure 4:**
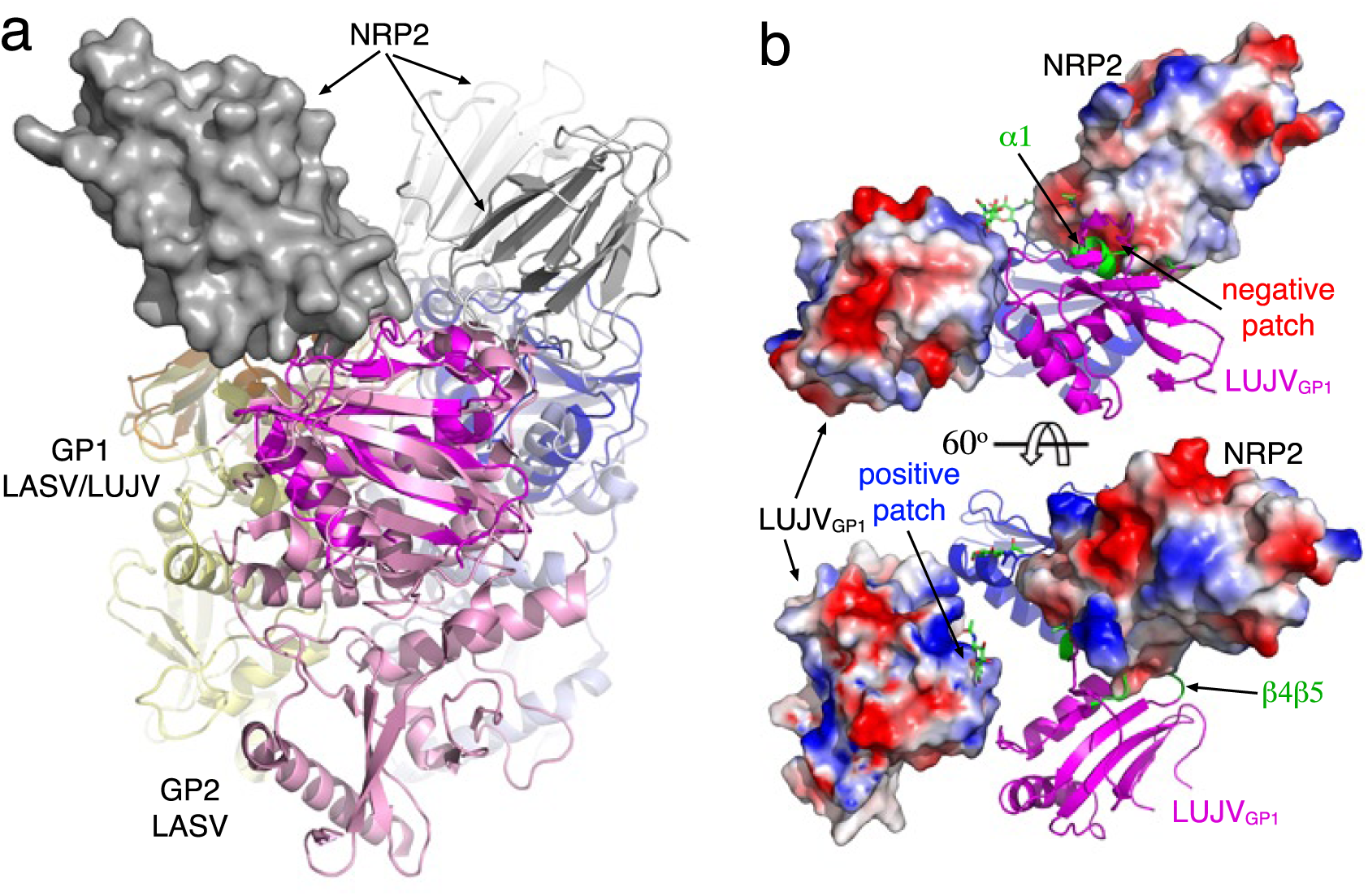
Binding to NRP2 involves a quaternary site that is composed of two copies of LUJV_GP1_ in the context of the spike. **a**. Three copies of LUJV_GP1_/NRP2 are superimposed on the three LASV_GP1_ copies in the structure of the trimeric spike^15^ (PDB: 5VK2). The three LUJV_GP1_ molecules are colored in orange, magenta, and blue, and the corresponding GP1/GP2 pairs from LASV are colored in yellow, pink, and light blue. The three copies of NRP2 are shown in gray using both ribbon and surface representations. The trimeric arrangement allows three NRP2 CUB domains to bind simultaneously. **b**, The quaternary binding site of NRP2. Three copies of LUJV_GP1_ in a trimeric configuration based on the superimposition in ‘a’ are shown with a single copy of NRP2. The local surface electrostatic potentials of NRP2 as well as one copy of LUJV_GP1_ are shown. The three copies of the first N-acetylglucosamine attached to the Asn112 residues are shown as green sticks. The α-helix 1 (α1) and β4β5 loop of LUJV_GP1_, which make the putative additional contacts with NRP2, are highlighted in green. The positive surface potential of α1, the β4β5 loop regions and the corresponding negatively charged region on NRP2 are marked. The lower view is tilted 60° with respect to the upper view.

Considering only the first CUB domain of NRP2, its binding angle is such that in principle up to three copies could be bound simultaneously without steric clashes (Fig. 4a). In this model, each NRP2 makes the contacts described above to one LUJV GP1 domains, but it also close to a neighboring LUJV_GP1_, potentially making additional contacts that are trimer dependent (Fig. 4b), as postulated for the α-DG binding of OW mammarenaviruses^15^ (Fig. 3b). In particular, NRP2 is in position to contact α-helix 1 and a loop connecting β-strands 4 and 5 (β4β5 loop) of the neighboring LUJV_GP1_. These contacts are near the glycan that is attached to Asn112 of LUJV_GP1_ (Fig. 1b and Fig. 4b). This glycan may restrict the access of antibodies to α-helix 1 and the β4β5 loop region as well as the β-strand 3 and α2β4 loop, which are nearby in the trimeric configuration. Although in this model the N-acetylglucosamine partially clashes with the NRP2 (Fig. 4b), a slight movement of the glycan toward the axis of the trimer will prevent such clashing. The positioning of LUJV_GP1_ in a trimeric configuration based on the spike complex of LASV is not accurate enough to elucidate the exact putative interactions that α-helix 1 and β4β5 loop may form with NRP2. Nevertheless, examining the contact electrostatic potential of NRP2 in the region predicted to form such interactions reveals a negatively charged patch (Fig. 4b). The surface that forms by α-helix 1 and β4β5 loop of LUJV_GP1_ is positively charged (Fig. 4b) and hence complementary. This observation supports the possibility that the complete NRP2 binding site forms by two neighboring LUJV_GP1_ molecules, helping to explain the fairly modest BSA upon complex formation that we observed in the crystal structure (Fig. 2a).

### Dissociation of LUJV_GP1_ from NRP2 in acidic conditions is likely due to changes in NRP2

The demonstration by Raaben *et al.* that LUJV_GP1_ dissociates from NRP2 at acidic pH^19^ raises the question whether pH-dependent binding could be explained at the structural level. Dissociation from NRP2 is important for efficient cell entry as it presumably allows LUJV to switch to CD63^19^ that may act as a triggering factor for membrane fusion in a similar way to LAMP1 in the case of LASV^17^. Since histidine has a side-chain pKa of about 6, it is an obvious candidate for controlling pH-dependent protein-protein interactions. Of all the histidine residues in the LUJV_GP1_/NRP2 complex, only His81 on the surface of NRP2 is located at the interface of the complex (Fig. 5a). Interestingly, His81 is not involved in any obvious polar interaction but rather makes Van Der Waals interactions with an apolar pocket on LUJV_GP1_ that is flanked by positively charged residues (Fig. 2d and Fig. 5b). Potentially, a protonated and positively charged His81 could thus repel LUJV_GP1_, which may promote the dissociation of the proteins. However, based on our structural analysis, we propose an additional mechanism that may act in concert with His81 to break the interaction between LUJV_GP1_ and NRP2 during cell entry. Since the LUJV_GP1_ interaction with NRP2 depends on a calcium ion bound to the first CUB domain of NRP2 (Fig. 2b, 2c and Supplementary Fig. 5), losing this calcium ion during cell entry will cause LUJV_GP1_ to dissociate from NRP2. The fact that no calcium ion was observed bound in the previously determined structure of the full-length NRP2^20^ indicates a limited affinity to calcium. In the early endosomes the calcium concentration drops to the low micromolar range, which is ∼1000 fold less compared to the extracellular space^28^ where LUJV_GP1_ first attaches to NRP2. Together with the acidification that can help to protonate the negatively charge groups that coordinate the calcium ion (Fig. 2b), NRP2 may thus lose its calcium upon entering early endosomes and thus further promote the dissociation of LUJV.

Overall, the structural data that we provided elucidate that LUJV utilizes a binding site formed by β-strand 3 and α2β4 loop on one monomer and may further span α-helix 1 and β4β5 loop on an adjacent LUJV_GP1_ in the context of the trimer. We postulate that a monoclonal antibody that will target any of these sites on LUJV_GP1_ in the context of the trimeric spike would block binding to NRP2 and thus may neutralize the virus. Hence, we propose to focus efforts to elicit such antibodies as a way to provide potential therapeutics against LUJV.

**Figure 5:**
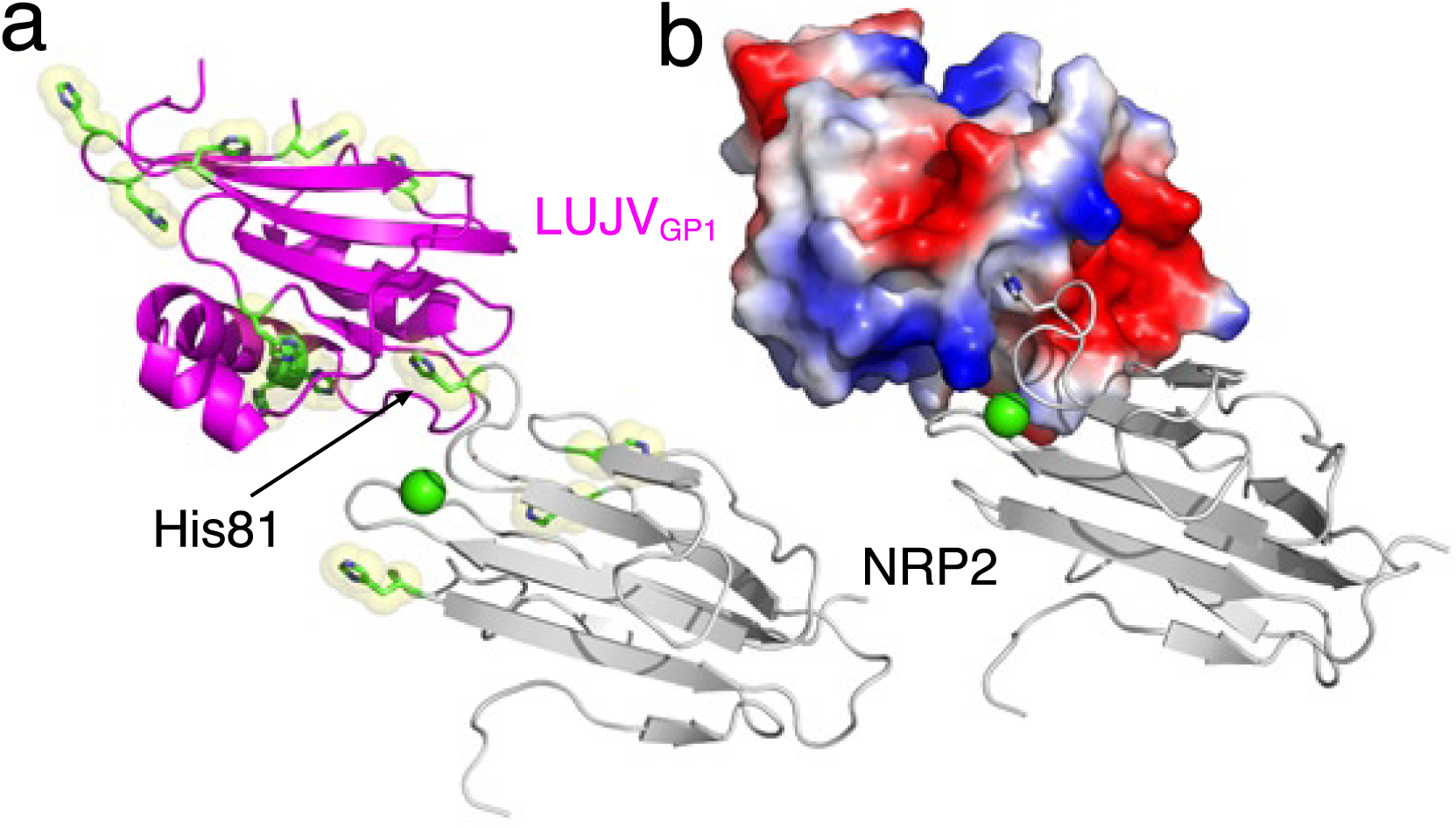
NRP2 contributes the sole histidine residue at the interface with LUJV_GP1_. **a**, A ribbon representation of the LUJV_GP1_/NRP2 complex. The histidine residues in both proteins are shown with green sticks and are highlighted using yellow space-filling spheres. The calcium ion is shown as a green sphere for reference. His81 at the interface is indicated. **b**, The contact surface potential of LUJV_GP1_ is shown. His81 of NRP2 is accommodated in an apolar pocket flanked by positively charged patches (blue surfaces).

## Methods

### Expression and purification of recombinant proteins

GP1_LUJV_ coding DNA (residues 74-199) was amplified from chemically synthesized LUJV GPC (Genscript). The gene was subcloned into the pACgp67b expression vector (BD Biosynthesis) to include an N-terminal 6x-His tag. Neuropilin 2 (NRP2) binding site fragment (residues 27-146) was chemically synthesized with C-terminal 6x-His tag, and was subcloned into the pACgp67b vector. Both GP1_LUJV_ and NRP2 were co-expressed as a secreted proteins using the baculovirus system as we previously described^11^. Cell media was collected, clarified using centrifugation and buffer exchanged to TBS (20 mM Tris-HCl pH 8.0, 150 mM sodium chloride) using a tangential flow filtration system (Millipore). LUJV_GP1_/NRP2 complex was captured using a HiTrap IMAC fast-flow Ni^+2^ (GE Healthcare) affinity column, and further purified using size exclusion chromatography with Superdex 75 10/300 column (GE Healthcare) in TBS buffer. Fc-fused NRP2 was expressed in HEK293 cells adapted to suspension cells (Expression Systems). Transfections were done using linear 25 kDa polyethylenimine (PEI) (Polysciences) at 1 mg of plasmid DNA per 1 L of culture at cell density of 1 M/ml. Media were collected after 5 days of incubation and supplemented with 0.02% (wt/vol) sodium azide and PMSF. Fusion proteins were isolated using protein-A affinity chromatography (GE Healthcare).

### Crystallization

We used vapor diffusion in sitting drops method for crystallization screens. For crystallization experiments we used a Mosquito® crystallization robot (TTP labs) to set 60, 120, and 180 nL drops of protein with 120 nL reservoir of commercially available crystallization screens. Initial crystallization hits for LUJV_GP1_/NRP2 were identified using PEGRx HT™ (Hampton) screen. Crystallization conditions were manually optimized. We further identified using additive screen HT™ (Hampton) the tri-peptide glycyl-glycyl-glycine as an additive that improves crystals’ morphology. Crystals for diffraction experiments were obtained by mixing the LUJV_GP1_/NRP2 complex solution at 12 mg/mL with 0.03M glycyl-glycyl-glycine, 25.9% PEG 6000, 0.09M Bis-Tris Propane pH 9.5 at 20° C. Crystals were briefly soaked in mother liquor solution supplemented with 25 % ethylene glycol for cryo preservation before flash cooling in liquid nitrogen.

### Data collection, structure solution and refinement

X-ray diffraction data were collected at the European Synchrotron Radiation Facility (ESRF) at beamline ID23-2 using a Pilatus 3 2M detector. Diffraction data were collected to a resolution of 2.4 Å. Images were indexed, integrated, and scaled using Xia2^29^ pipeline that made use of aimless^30^, CCP4^31^, Dials^32^, and Pointless^33^. A molecular replacement solution using the first CUB domain of NRP2^20^ (PDB: 2QQK) was found using Phaser^21^. We used Phenix.AutoBuild^22^ to start tracing LUJV_GP1_ and subsequently extend the model to completion using manual building in Coot^23^.

### Cell stain

HEK293T cells were seeded on poly-L-Lysine pre-coated cover slips in 24-well plates and transfected with a plasmid (pcDNA3) encoding LUJV GPC using PEI-MAX (polysciences) reagent. At 48 h post transfection cells were fixed with pre-warmed 3.7% formaldehyde (PFA) solution in PBS and blocked with 3% BSA in PBS. Cover slips were incubated with NRP2-Fc diluted in 1% BSA-PBS at a concentration of 30 μg/ml, with or without addition of 5 mM EGTA (Sigma), followed by staining with Cy3-conjugated anti-Human Fc (Jackson) and mounting with mounting media supplemented with DAPI (GBI Labs). Cells were imaged at ×10 magnification and images were processed using ImageJ^34^.

### Accession code

Atomic model for the LUJV_GP1_/NRP2 complex as well as structure factors were deposited to the protein data bank under accession code 6GH8.

## Acknowledgments

Diffraction experiments were performed in beamline ID23-2 at the European Synchrotron Radiation Facility (ESRF), Grenoble, France. We are grateful to Chloe Zubieta at the ESRF for providing assistance in using beamline ID23-2. We thank professor Randy Read from University of Cambridge for his invaluable advises and contribution in analyzing our crystallographic data. We thank professor Deborah Fass for providing critical comments and suggestions. Ron Diskin is incumbent of the Tauro career development chair in biomedical research. Research in the Diskin lab is supported by a research grant from the Enoch Foundation, a research grant from the Abramson Family Center for Young Scientists, a research grant from Ms. Rudolfine Steindling, by the Minerva Foundation with funding from the Federal German Ministry for Education and Research, and by a grant from the Israel Science Foundation (grant No. 682/16).

## Author contributions

H.C.D. together with I.K. produced, purified and crystallized the LUJV_GP1_/NRP2 complex. H.C.D. and R.D. collected diffraction data. R.D. solved and analyzed the structure and wrote the manuscript with the help of H.C.D and I.K.

## References

1 Moraz, M. L. & Kunz, S. Pathogenesis of arenavirus hemorrhagic fevers. Expert Rev Anti Infect Ther 9, 49–59, doi:10.1586/eri.10.142 (2011).

2 Geisbert, T. W. & Jahrling, P. B. Exotic emerging viral diseases: progress and challenges. Nat Med 10, S110–121, doi:10.1038/nm1142 (2004).

3 Nunberg, J. H. & York, J. The curious case of arenavirus entry, and its inhibition. Viruses 4, 83–101, doi:10.3390/v4010083 (2012).

4 Cao, W. et al. Identification of alpha-dystroglycan as a receptor for lymphocytic choriomeningitis virus and Lassa fever virus. Science 282, 2079–2081 (1998).

5 Kunz, S., Rojek, J. M., Perez, M., Spiropoulou, C. F. & Oldstone, M. B. Characterization of the interaction of lassa fever virus with its cellular receptor alpha-dystroglycan. J Virol 79, 5979–5987, doi:10.1128/JVI.79.10.5979-5987.2005 (2005).

6 Radoshitzky, S. R. et al. Transferrin receptor 1 is a cellular receptor for New World haemorrhagic fever arenaviruses. Nature 446, 92–96, doi:10.1038/nature05539 (2007).

7 Flanagan, M. L. et al. New world clade B arenaviruses can use transferrin receptor 1 (TfR1)-dependent and -independent entry pathways, and glycoproteins from human pathogenic strains are associated with the use of TfR1. J Virol 82, 938–948, doi:10.1128/JVI.01397-07 (2008).

8 Eschli, B. et al. Identification of an N-terminal trimeric coiled-coil core within arenavirus glycoprotein 2 permits assignment to class I viral fusion proteins. J Virol 80, 5897–5907, doi:10.1128/JVI.00008-06 (2006).

9 Rojek, J. M. & Kunz, S. Cell entry by human pathogenic arenaviruses. Cellular microbiology 10, 828–835, doi:10.1111/j.1462-5822.2007.01113.x (2008).

10 Bowden, T. A. et al. Unusual molecular architecture of the machupo virus attachment glycoprotein. J Virol 83, 8259–8265, doi:10.1128/JVI.00761-09 (2009).

11 Cohen-Dvashi, H., Cohen, N., Israeli, H. & Diskin, R. Molecular Mechanism for LAMP1 Recognition by Lassa Virus. J Virol 89, 7584–7592, doi:10.1128/JVI.00651-15 (2015).

12 Israeli, H., Cohen-Dvashi, H., Shulman, A., Shimon, A. & Diskin, R. Mapping of the Lassa virus LAMP1 binding site reveals unique determinants not shared by other old world arenaviruses. PLoS Pathog 13, e1006337, doi:10.1371/journal.ppat.1006337 (2017).

13 Shimon, A., Shani, O. & Diskin, R. Structural basis for receptor selectivity by the Whitewater Arroyo mammarenavirus. J Mol Biol, doi:10.1016/j.jmb.2017.07.011 (2017).

14 Mahmutovic, S. et al. Molecular Basis for Antibody-Mediated Neutralization of New World Hemorrhagic Fever Mammarenaviruses. Cell Host Microbe 18, 705–713, doi:10.1016/j.chom.2015.11.005 (2015).

15 Hastie, K. M. et al. Structural basis for antibody-mediated neutralization of Lassa virus. Science 356, 923–928, doi:10.1126/science.aam7260 (2017).

16 Jae, L. T. et al. Virus entry. Lassa virus entry requires a trigger-induced receptor switch. Science 344, 1506–1510, doi:10.1126/science.1252480 (2014).

17 Cohen-Dvashi, H., Israeli, H., Shani, O., Katz, A. & Diskin, R. Role of LAMP1 Binding and pH Sensing by the Spike Complex of Lassa Virus. J Virol 90, 10329–10338, doi:10.1128/JVI.01624-16 (2016).

18 Briese, T. et al. Genetic detection and characterization of Lujo virus, a new hemorrhagic fever-associated arenavirus from southern Africa. PLoS Pathog 5, e1000455, doi:10.1371/journal.ppat.1000455 (2009).

19 Raaben, M. et al. NRP2 and CD63 Are Host Factors for Lujo Virus Cell Entry. Cell Host Microbe 22, 688–696 e685, doi:10.1016/j.chom.2017.10.002 (2017).

20 Appleton, B. A. et al. Structural studies of neuropilin/antibody complexes provide insights into semaphorin and VEGF binding. EMBO J 26, 4902– 4912, doi:10.1038/sj.emboj.7601906 (2007).

21 McCoy, A. J. et al. Phaser crystallographic software. J Appl Crystallogr 40, 658–674, doi:10.1107/S0021889807021206 (2007).

22 Adams, P. D. et al. PHENIX: a comprehensive Python-based system for macromolecular structure solution. Acta crystallographica. Section D, Biological crystallography 66, 213–221, doi:10.1107/S0907444909052925 (2010).

23 Emsley, P., Lohkamp, B., Scott, W. G. & Cowtan, K. Features and development of Coot. Acta crystallographica. Section D, Biological crystallography 66, 486–501, doi:10.1107/S0907444910007493 (2010).

24 Hastie, K. M. et al. Crystal structure of the prefusion surface glycoprotein of the prototypic arenavirus LCMV. Nat Struct Mol Biol 23, 513–521, doi:10.1038/nsmb.3210 (2016).

25 Abraham, J., Corbett, K. D., Farzan, M., Choe, H. & Harrison, S. C. Structural basis for receptor recognition by New World hemorrhagic fever arenaviruses. Nat Struct Mol Biol 17, 438–444, doi:10.1038/nsmb.1772 (2010).

26 Sullivan, B. M. et al. Point mutation in the glycoprotein of lymphocytic choriomeningitis virus is necessary for receptor binding, dendritic cell infection, and long-term persistence. Proc Natl Acad Sci U S A 108, 2969– 2974, doi:10.1073/pnas.1019304108 (2011).

27 Smelt, S. C. et al. Differences in affinity of binding of lymphocytic choriomeningitis virus strains to the cellular receptor alpha-dystroglycan correlate with viral tropism and disease kinetics. J Virol 75, 448–457, doi:10.1128/JVI.75.1.448-457.2001 (2001).

28 Scott, C. C. & Gruenberg, J. Ion flux and the function of endosomes and lysosomes: pH is just the start: the flux of ions across endosomal membranes influences endosome function not only through regulation of the luminal pH. Bioessays 33, 103–110, doi:10.1002/bies.201000108 (2011).

29 Winter, G. xia2: an expert system for macromolecular crystallography data reduction. Journal of Applied Crystallography 43, 186–190, doi:doi:10.1107/S0021889809045701 (2010).

30 Evans, P. R. & Murshudov, G. N. How good are my data and what is the resolution? Acta crystallographica. Section D, Biological crystallography 69, 1204–1214, doi:10.1107/S0907444913000061 (2013).

31 Winn, M. D. et al. Overview of the CCP4 suite and current developments. Acta crystallographica. Section D, Biological crystallography 67, 235–242, doi:10.1107/S0907444910045749 (2011).

32 Winter, G. et al. DIALS: implementation and evaluation of a new integration package. Acta Crystallogr D Struct Biol 74, 85–97, doi:10.1107/S2059798317017235 (2018).

33 Evans, P. Scaling and assessment of data quality. Acta crystallographica. Section D, Biological crystallography 62, 72–82, doi:10.1107/S0907444905036693 (2006).

34 Schneider, C. A., Rasband, W. S. & Eliceiri, K. W. NIH Image to ImageJ: 25 years of image analysis. Nat Methods 9, 671–675 (2012).

